# A Genetic Algorithm to scour protein sequence space: A novel framework for protein engineering using protein language models and force fields

**DOI:** 10.1101/2025.10.29.685431

**Authors:** João Sartori, Eduardo Krempser, Ana Carolina Ramos Guimarães, Lucas de Almeida Machado

**Author notes:** Correspondence should be addressed to Guimarães, A. C. R.

## Abstract

The optimization of protein sequences for enhanced binding and stability remains a formidable challenge in bioengineering due to the vastness of sequence space. Existing state-of-the-art methods, including traditional structure-based design and protein language models, use fitness estimators as objective functions to guide search algorithms that scour sequence space for optimal sequences. However, effective exploration requires search strategies that balance diversity and computational efficiency, as both are paramount for effective exploration. This work presents GAPO (Genetic Algorithm for Protein Optimization), a novel flexible framework that integrates evolutionary computing, protein language models, and structure-based design to efficiently explore sequence space. GAPO employs genetic algorithms to iteratively refine protein sequences based on user-defined objective functions, using either force field-derived information or protein language models to assess fitness, it also allows users to define custom objective functions, including multi-objective ones. We detail GAPO’s implementation, highlighting features such as customizable initialization methods, diverse selection strategies, and mutation techniques informed by evolutionary scale modeling (ESM).IIn a case study using Hen-egg lysozyme, GAPO outperformed simulated annealing (SA) in both protein language model and energy-based objectives, converging to higher average ESM2 probabilities (0.98 vs. WT 0.89 and SA 0.88) and more favorable REF2015 energies (−510 REU vs. WT −415 REU and SA −405 REU), while maintaining reproducible behavior across independent runs. GAPO is available at https://github.com/izzetbiophysicist/GAPO.

## Introduction

The exploration of protein sequence space for the discovery of optimal binders, catalysts or more stable structures represents one of the most significant challenges in bioengineering. The vastness of this space, encompassing all possible amino acid permutations for a given protein length, makes exhaustive search impractical (Vila 2020), or virtually impossible . As a result, sophisticated in silico search methods have become indispensable in guiding the search toward sequences with desirable properties (Tretyachenko et al. 2021). Traditional structure-based protein design approaches, such as those implemented in the Rosetta suite (Leaver-Fay et al. 2011), leverage force field-based methods to predict stability, binding affinity and other features. These techniques have been instrumental in designing proteins with enhanced stability, affinity, and specificity(Tinberg et al. 2013; Sitthiyotha and Chunsrivirot 2021; Warszawski et al. 2019). In contrast, sequence-based design strategies, such as consensus design, capitalize on evolutionary information to identify conserved residues that contribute to stability or function. More recently, the advent of artificial intelligence (AI) methods, particularly large language models such as ESM (Evolutionary Scale Modeling) (Lin et al. 2023), has introduced a new paradigm in protein engineering, being able to consider evolutionary constraints as conditional probabilities. These models predict sequence-function relationships with remarkable accuracy by capturing the statistical properties of natural sequences (Yan Wang et al. 2025) . Despite their promise, AI-driven approaches often suffer from a lack of efficient search mechanisms, relying on simple search strategies (Hie et al. 2024) or methods such as Beam search (Ferruz, Schmidt, and Höcker 2022) or greedy strategies using point mutations (Sun, Cui, and Wu 2021) which limits their applicability in exploring a wider breadth of sequence space, mainly because it can be difficult to optimize the sequences taking into consideration a large enough array of conditional probabilities and because many of these methods start from a single sequence and expand by iterative modifications. Recent studies also attempted merging ESM and Rosetta-based protein design, enhancing the Protein Language Model (PLM) scores of the protein sequences (Ertelt, Meiler, and Schoeder 2024), but the search problems are not absent here either, if anything, the search space can become even more cumbersome when considering the many ways by which force field-derived and PLM scores can be combined.

Evolutionary computing offers a robust solution to this problem, particularly through the application of genetic algorithms (GAs). GAs are a class of optimization algorithms inspired by the principles of natural selection and genetic inheritance (Holland 2019). They are particularly well-suited for navigating complex, high-dimensional search spaces, making them ideal for protein engineering tasks where the objective is to optimize specific properties, such as binding affinity or stability, across a vast sequence landscape.

This manuscript introduces GAPO (Genetic Algorithm for Protein Optimization), a novel software that integrates the strengths of genetic algorithms with the predictive power of structure-based and sequence-based design methods. GAPO provides an efficient search mechanism for identifying optimal protein sequences, offering a versatile toolkit for the next generation of protein engineering challenges. Here we show that GAPO consistently beats Simulated Annealing in both force field-guided search and ESM2 guided search in our case study.

## Implementation

GAPO’s framework is designed to streamline the process of protein optimization by employing a genetic algorithm that iteratively improves protein sequences based on user-defined objective functions and parameters. The workflow of GAPO is structured to allow flexibility and customization, ensuring that it can be adapted to a wide range of protein engineering tasks. The following sections are a detailed description of the methodology employed by GAPO and each of its components.

GAPO was fully implemented in Python, version 3.7, and besides the standard packages it depends on ESM2 (Lin et al. 2023), and pyRosetta4(Chaudhury, Lyskov, and Gray 2010) in order to work. The main script (GAprot.py) imports an assortment of language model and structure manipulation functions from “apt_functions.py”, which also contains a series of predefined fitness functions. The main script runs all the populations inside a “GeneticAlgoBase” class, which contains all the parameters and information of the proteins. A third script is used to import GA_main.py and run the optimization.

### Initialization

The process begins with the generation of an initial population of protein sequences. This population can be generated in various ways depending on the specific goals of the optimization:

i. Random Initialization: Sequences are randomly generated, providing a broad exploration of the sequence space.
ii. Seeded Initialization: The population is seeded with known sequences, such as natural proteins or engineered variants, to focus the search on regions of sequence space for which the user has strong prior knowledge about protein fitness.

Each sequence within the population is treated as an individual in the context of the GA, and the initial fitness of each individual is determined by the evaluation of user-defined objective functions. In all its instances, GAPO only works with populations of sequences that have the same length. If structure-based optimization is being used, a reference structure should be provided, then the sequences will be modeled using the reference structure and pyRosetta’s repacking and fast_relax routines. During this step, the user can select the residues he wants to fixate during the optimization.

### Fitness Evaluation

Fitness evaluation is a critical step in the genetic algorithm, as it guides the selection process. GAPO offers a diverse array of objective functions through the apt_function module, allowing users to assess sequences based on multiple criteria.

i. Structural Stability: Evaluation based on force field calculations using pyRosetta.
ii. Average ESM probability: Average residue probability as predicted by ESM2.

These fitness functions can be used individually or in combination, which can be easily implemented by advanced users, with weighted sums allowing for multi-objective optimization. This flexibility enables the fine-tuning of the optimization process to target specific protein properties.

### Selection

The selection process determines which sequences will contribute to the next generation. GAPO supports various selection strategies, each with its advantages:

i. Tournament Selection: Randomly chosen subsets of the population are pitted against each other, with the best-performing sequences selected for reproduction. Tournament sizes and rounds can be set by the user.
ii. Elitism: A strategy that ensures the top-performing sequences are always carried over to the next generation, preventing loss of the best solutions.

By combining these strategies, GAPO maintains a balance between exploration (diversifying the sequence space) and exploitation (refining promising sequences).

### Crossover

Crossover is a genetic operation that recombines two parent sequences to generate offspring, mimicking the process of recombination in natural evolution. The crossover operation is critical for introducing new sequence variations that may lead to improved fitness in subsequent generations (Figure 1). In GAPO, crossover can be performed in various ways:

**Figure 1:**
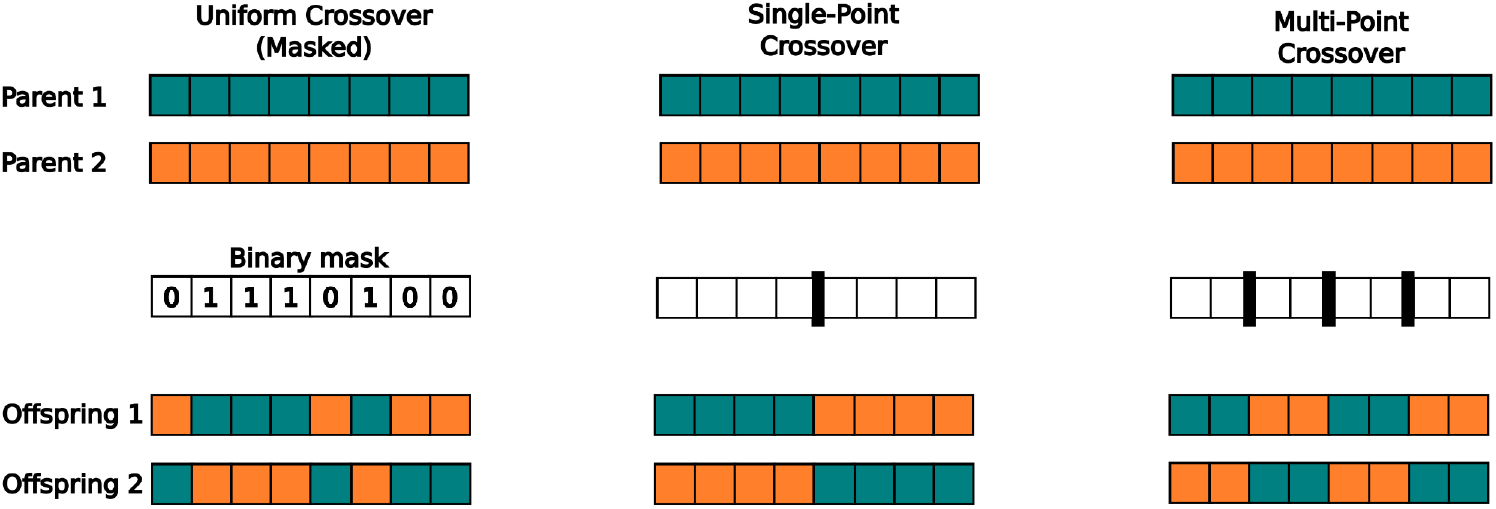
Schematic illustration of three distinct crossover operators in genetic algorithms.

i. Uniform Crossover (masked crossover): Each position in the offspring sequence is randomly selected from one of two parents, creating a more uniform blend of the parental sequences.
ii. Single-Point Crossover: A single crossover point is chosen along the sequence, and the segments before and after the point are swapped between the two parent sequences. The crossover can always be carried at the midpoint between the protein sequences, or may be set to fluctuate around the middle by an adjustable degree.
iii. Multi-Point Crossover: Multiple crossover points are selected, resulting in more complex recombinations that can introduce greater diversity into the population.

### Mutation

After each crossover it will be decided if the sequence is mutated using the mutation rate, and then a random position is mutated, further enhancing the diversity of the population. It ensures that the population does not converge prematurely and continues to explore new areas of the sequence space. GAPO allows users to control the mutation rate and type. The mutations can be completely random or can be added using ESM2, to raise the likelihood of mutating the position to a residue that’s likely to occur given the rest of the sequence. The temperature of the ESM2-based inference can be chosen by the user.

## Case study: Optimizing Hen-Egg lysozyme’s stability

As a case study to test the capability of GAPO to optimize ESM2 likelihoods or REF15 forcefield scores, we attempted optimizing the structural stability of Hen-Egg Lysozyme (PDB: 2LZT) (Ramanadham, Sieker, and Jensen 1990) employing two distinct strategies: a sequence-driven and a force field-driven approach.

### Seeded initialization

Each optimization strategy was initiated from one of two distinct seed populations, prepared as follows:

i. Natural Sequence Seeding: A pool of sequences was retrieved from the UniProtKB/Swiss-Prot database (queried August 9, 2024) (UniProt Consortium, 2023) using the term “lysozyme” and filtering for reviewed entries. These sequences were then filtered, retaining only those with >35% sequence identity and >90% query coverage relative to the reference (2LZT). To form a diverse starting population of 200, a pairwise distance matrix was calculated using the BLOSUM62 matrix (Henikoff and Henikoff 1992). A Monte Carlo sampling of 10,000 subsets was performed, and the subset with the maximum cumulative pairwise distance was selected.
ii. ESM2-based Seeding: The sequences were generated using the esm2_t33_650M_UR50D. The wild-type (WT) sequence of 2LZT served as the initial prompt, and 200 new sequences were generated using a sampling temperature of 1.5 to encourage novelty.

### Sequence Standardization and Optimization Constraints

All sequences, regardless of their origin, were first standardized by aligning them to the reference sequence and grafting them onto the 2LZT reference sequence. This ensured uniform length and structural compatibility for all variants before optimization.

To preserve essential functions, specific residues were constrained (locked) from mutation during the optimization runs, including all cysteine residues involved in structurally critical disulfide bonds, and residues forming the active site and substrate-binding pocket. This set was defined by superimposing the 2LZT structure onto PDB entry 2DQA (a lysozyme-substrate complex) (Goto et al. 2007) and selecting all residues within a 6 Å radius of the bound substrate.

### Optimization

The optimization process employed tournament selection, a method chosen for its balance between exploration and exploitation of the sequence space. The selection involved 50 tournament cycles, with each cycle comprising 4 individuals per tournament. This approach allowed for a focused refinement of the population while maintaining a diverse pool of candidates.

The optimization was conducted over 100 cycles, iteratively improving the population’s fitness. Mutations were introduced at a rate of 90%, using the ESM2 to predict residue substitutions. The elevated mutation rate is justified by the use of ESM2, which has strong priors about the residue fitness (Hie et al. 2024). This approach ensured that mutations were contextually appropriate, preserving the overall structure and function of lysozyme. In structure-driven optimization, fitness evaluation was performed using pyRosetta’s REF15 energy function, which served as the objective measure of structural stability. The REF15 energy function allowed for the accurate assessment of the lysozyme’s stability, guiding the genetic algorithm toward sequences that exhibited enhanced stability. In sequence-driven optimization we used only ESM2 average sequence probability as fitness function and no structures were used. The different parameters combinations are referred to by fitness function - Fitness(Rosetta) or Fitness(ESM), respectively using REF15 or ESM2 average sequence probability -; and by mutation method - Mut(ESM) or Mut(Random), respectively using ESM2 residue probabilities or an uniform probability distribution.

As a control we used Simulated annealing (SA) runs for each sequence of the starting population from the ESM2 temperature 1.5. Each SA trajectory consisted of 100 iterations, where in each step a single point mutation was introduced. The mutation position was selected uniformly at random, and the new amino acid was proposed by the esm2_t33_650M_UR50D model.

### Structure analysis

For structural validation, the top-performing variant from each optimization condition and the WT sequence were selected. Their 3D structures were predicted using the AlphaFold3 (Abramson et al. 2024) server with default parameters and subsequently refined with three cycles of the PyRosetta FastRelax protocol(Chaudhury, Lyskov, and Gray 2010). A pairwise non-bonded energy matrix was then calculated for each refined structure by summing the fa_atr, fa_rep, fa_elec, fa_sol, and lk_ball_wtd terms from the REF2015 score function (Alford et al. 2017). To focus on physically relevant interactions, this matrix was filtered to include only pairs of residues within a 6.0 Å C-alpha distance. The final energetic network was visualized in PyMOL as CGO cylinders, representing the most significant interactions, with colors indicating more negative favorable (blue) or more positive (red) energies.

## Results

Across all experimental conditions, the GA optimized the target metrics, whether focusing on the average ESM2 probability or improving the ΔGfold of the lysozyme. All optimization regimens with GA were capable of obtaining fitness measures that are, as expected, better than the WT enzyme (Figures 2).

**Figure 2:**
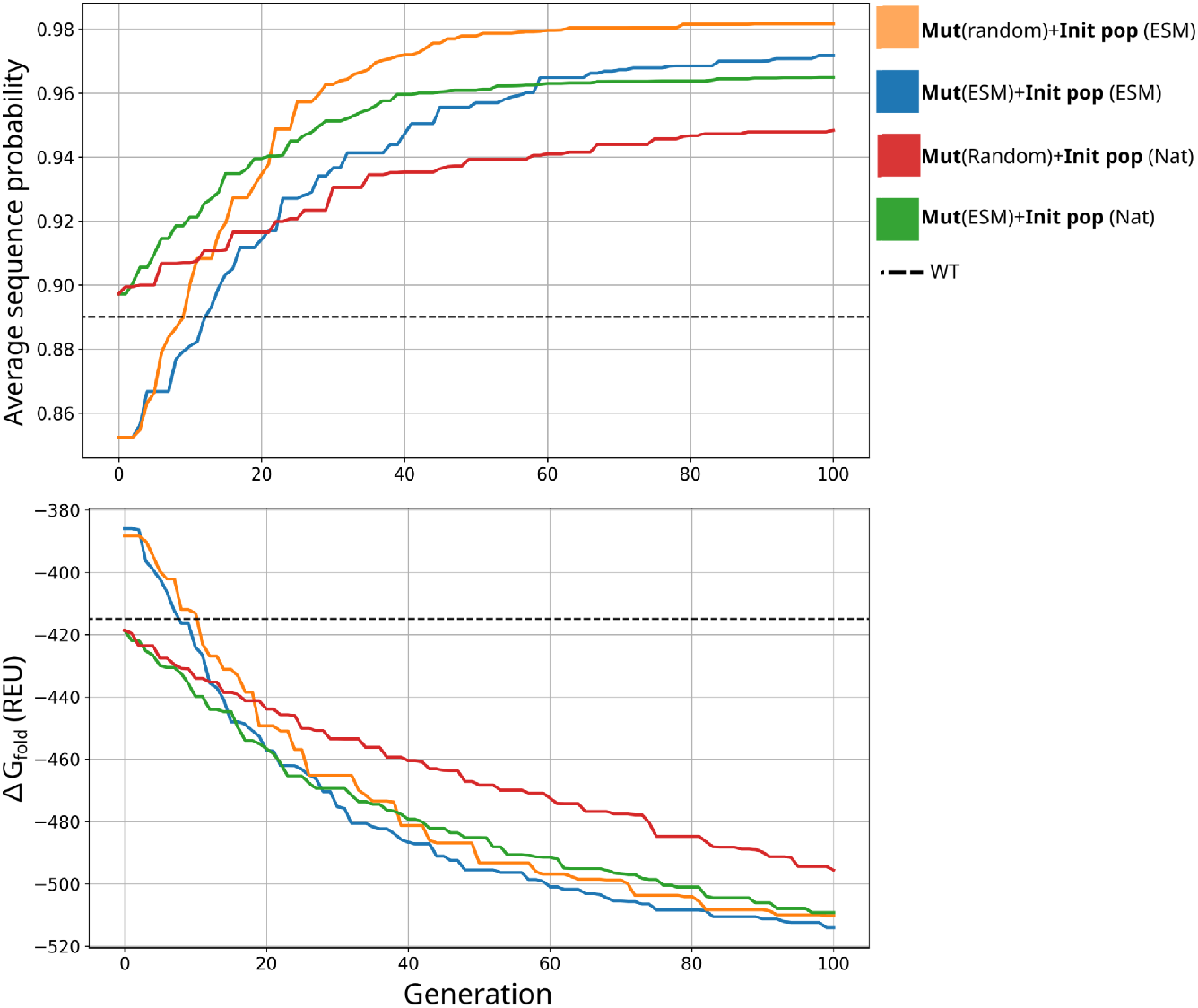
Evolution of the best individual per generation using the natural sequences as the initial population. a) Trajectory of Average sequence probability of each generation’s best individual when using ESM2 probability as fitness function; b) trajectory of the REF15 ΔGfold of each generation’s best individual when using REF15 as fitness function. The combination of parameters are shown as blue, orange lines, green and red, the black dashed line represents the reference value of the wild-type protein.

While optimizations initiated with natural (NAT) sequences started from a higher fitness baseline, those starting from ESM-generated populations consistently converged to superior final optimization scores. This outcome holds true for both the ESM2 and Rosetta fitness functions, suggesting that the diversity within the ESM2-generated ensembles provides a more fertile landscape for optimization, despite a lower initial fitness.

Regarding the mutation method, the guided approach using ESM2 to introduce mutations consistently outperformed random mutations. This was observed for both the NAT and ESM-derived starting populations across both fitness functions, indicating that leveraging the model’s learned representations is a more effective strategy for navigating the sequence space toward highly optimized variants, even when the fitness function uses a molecular forcefield.

Notably, the effectiveness of ESM-guided mutations in optimizing a physics-based metric such as Rosetta’s REF15 highlights the relevance of PLMs beyond sequence-based fitness prediction. This synergy suggests that PLMs capture abstract features of protein fitness that are complementary to physics-based energy functions (Ertelt, Meiler, and Schoeder 2024). These findings, therefore, indicate a promising avenue for future research on hybrid methodologies. Indeed, this prospect is directly enabled by our framework, as GAPO allows users to combine fitness functions. This capability would permit the optimization of a system using both ESM2 and REF15 concurrently, thereby creating a novel, composite fitness landscape that capitalizes on the strengths of both data-driven and physics-based approaches. To validate GAPO’s performance against established optimization methods, we benchmarked it against the SA algorithm (Figure 3). Both methods were employed to optimize sequences using the ESM2 score and the REF15 energy as objective functions, with all mutations guided by ESM throughout the experiments.

**Figure 3:**
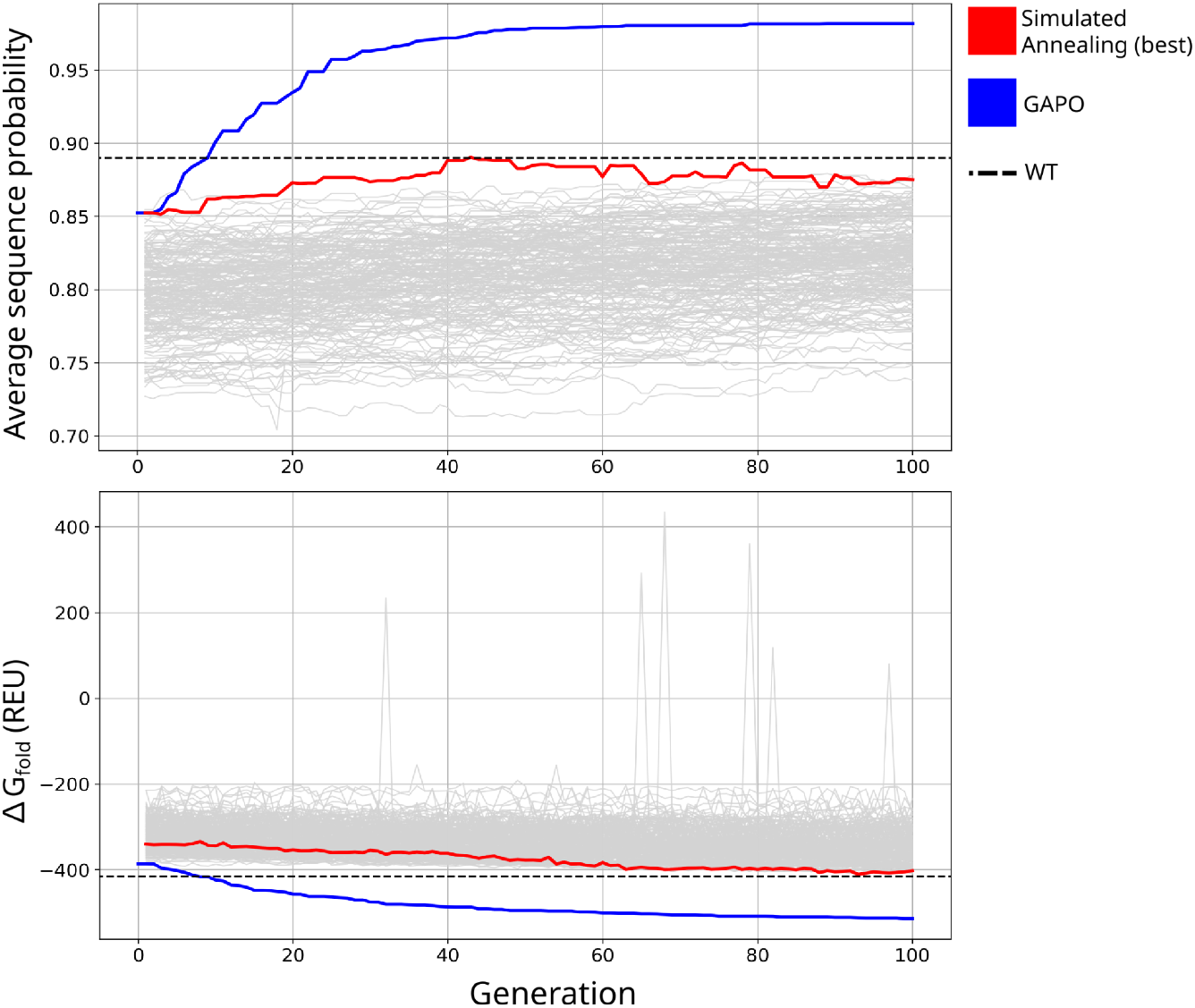
Benchmark comparing the evolution of the best individual from each generation using simulated annealing and GAPO. a) Trajectory of Average sequence probability of each generation’s best individual when using ESM2 probability as fitness function; b) trajectory of the REF15 ΔGfold of each generation’s best individual when using REF15 as fitness function. The combination of parameters are shown as blue and red, the black dashed line represents the reference value of the wild-type protein.

The results consistently demonstrate GAPO’s superior performance. When optimizing the ESM2 score, GAPO rapidly converged to a final score of approximately 0.98, significantly surpassing the WT baseline of 0.89. In contrast, Simulated Annealing achieved a final score of approximately 0.88, failing to outperform the WT baseline. GAPO’s advantage was even more pronounced during the REF15 energy optimization, where it reached a final energy value of approximately −510 REU, representing a substantial improvement over the WT score of −415 REU. Conversely, the SA optimization concluded at an energy of approximately −405 REU, unable to reach the WT energy level.

To investigate the diversity of all the generated populations, we analyzed the sequence identity of the generated sequences relative to the wild-type (WT) sequence for each optimization condition (Figure 4).

**Figure 4:**
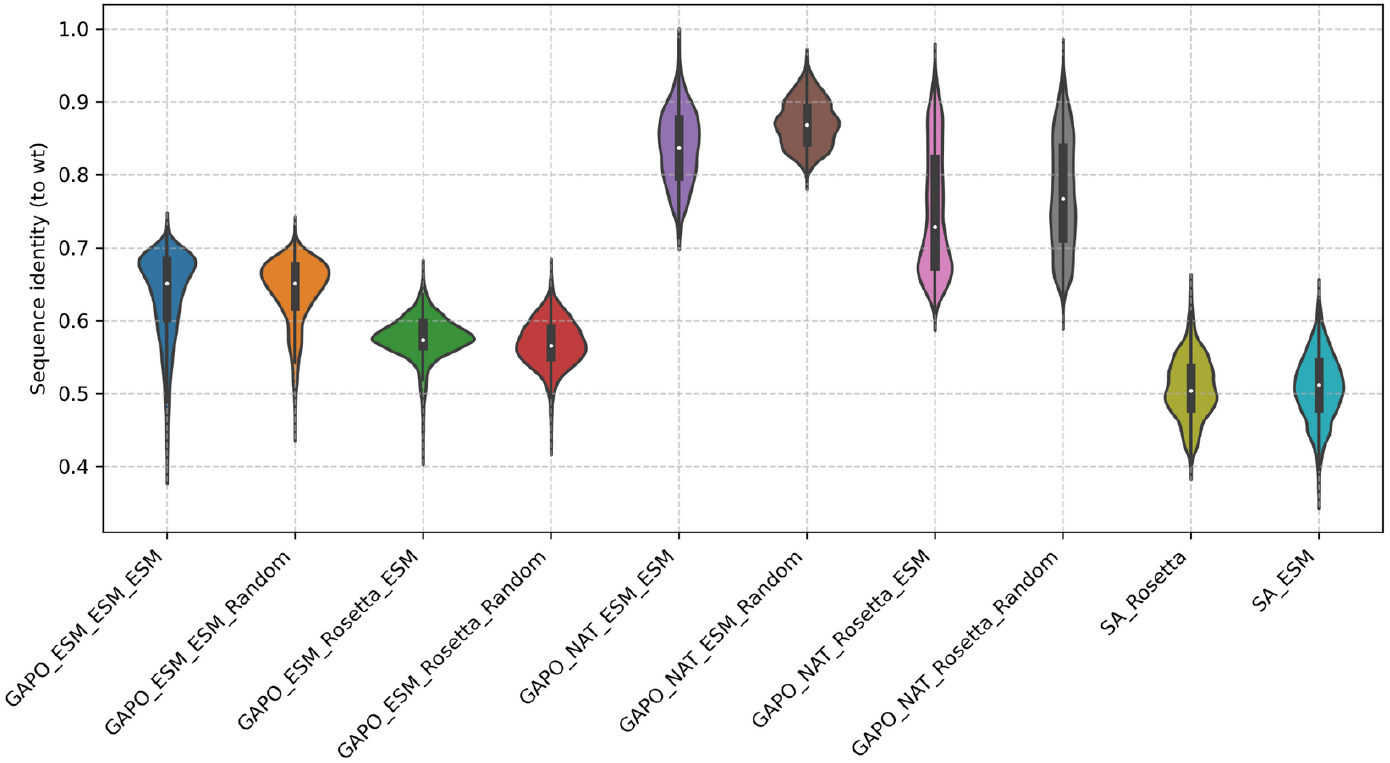
Sequence identity distribution of the final populations. Violin plots showing the sequence identity of each generated variant relative to the wild-type (WT) sequence. Each violin represents the population’s distribution for a specific optimization condition, varying the algorithm (GAPO or SA), initial population (ESM or NAT), fitness function (ESM or Rosetta), and mutation method (ESM or Random).

The resulting distributions reveal a clear impact between the optimization strategy and the sequence diversity when looking at the initialization and optimization strategy. Optimizations initiated from natural sequences (GAPO_NAT_*) consistently produced populations with lower diversity, evidenced by a higher median sequence identity. This suggests these trajectories explored a more constrained region of the sequence space around the starting point. Conversely, the SA baseline method, despite its suboptimal performance in fitness optimization, generated the most diverse ensembles, exhibiting the lowest median sequence identity, which can be explained by the fact that it has multiple independent runs that can take different directions. This indicates that while the SA strategy promotes a broad exploration of the sequence space, this exploratory capacity did not effectively translate into convergence towards high-fitness solutions in our experiments.

To provide a more granular view of sequence diversity and convergence beyond average identity, we mapped the optimization trajectories using PLMs embeddings. We generated ESM2 embeddings for all sequences and used Principal Component Analysis (PCA) to visualize their distribution (Figure 5).

**Figure 5:**
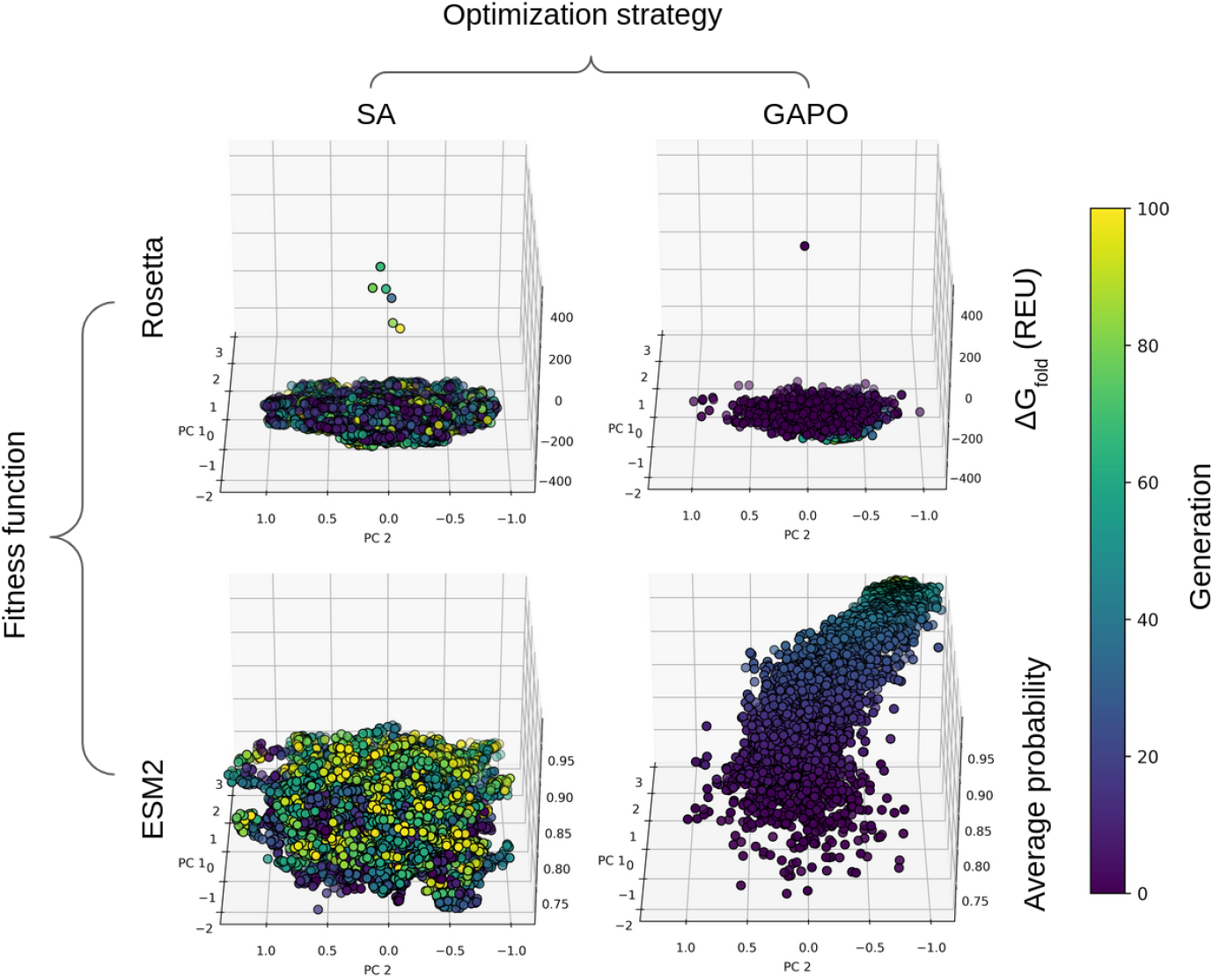
Sequence diversity throughout the optimization using ESM2-generated sequences as the initial population. Each point represents a protein sequence, with its position determined by the first two principal components (explaining 71% and 9% of the variance, respectively) of its ESM2 embedding. Rows correspond to different fitness functions: the top row shows the optimization of REF15 scores, while the bottom row shows the optimization of the average residue probability from ESM2. Columns represent the optimization strategy: Simulated Annealing on the left and GAPO on the right. Points are color-coded by generation, progressing from early generations (purple) to the latest (yellow).

While the SA strategy produced ensembles with greater sequence diversity relative to the wild-type (as shown in Figure 5), the PCA of the embedding space reveals that this exploration is largely undirected. For both Rosetta and ESM2 fitness functions, the SA populations remain as a disorganized cloud, with later-generation variants (yellow) often failing to localize in regions of higher fitness. This suggests that SA performs a broad, stochastic search of the sequence space but struggles to effectively exploit promising regions to climb the fitness landscape. Its ‘diversity’ is therefore a sign of an inefficient search, not a productive exploration.

In contrast, GAPO demonstrates a clear convergent evolutionary trajectory. In all cases, the population visibly progresses from a diffuse starting point (purple) to a more condensed, high-fitness region (yellow). This convergence in the embedding space is direct evidence of effective optimization, where the algorithm successfully identifies and refines solutions within a promising area of the sequence landscape.

To understand the structural basis for the improved fitness scores, we modeled all of the best sequences using AlphaFold3 and analyzed their mean pLDDT and REU after relaxation rounds. Interestingly, SA_Rosetta and GAPO_Rosetta had REU values much higher than the WT enzyme, suggesting that REF15 as a metric - at least using single structures - can misguide the search; meanwhile, SA_ESM and GAPO_ESM had approximately an order or magnitude lower REU values when compared to their Rosetta-guided counterparts. It suggests that ESM2 is capable of leading towards a more stable route when compared to Rosetta, even though it does not consider physics in any explicit manner. We also used the structures to visualize the pairwise interaction networks of the top-performing variants and the wild-type (Figure 6). In this representation, each cylinder depicts a key pair-wise interaction between two residues, allowing for the intuitive identification of interaction hotspots. The color of the cylinders reveals the nature of the interaction: blue indicates a favorable (more negative) interaction, while red indicates an unfavorable (more positive) interaction.

**Figure 6:**
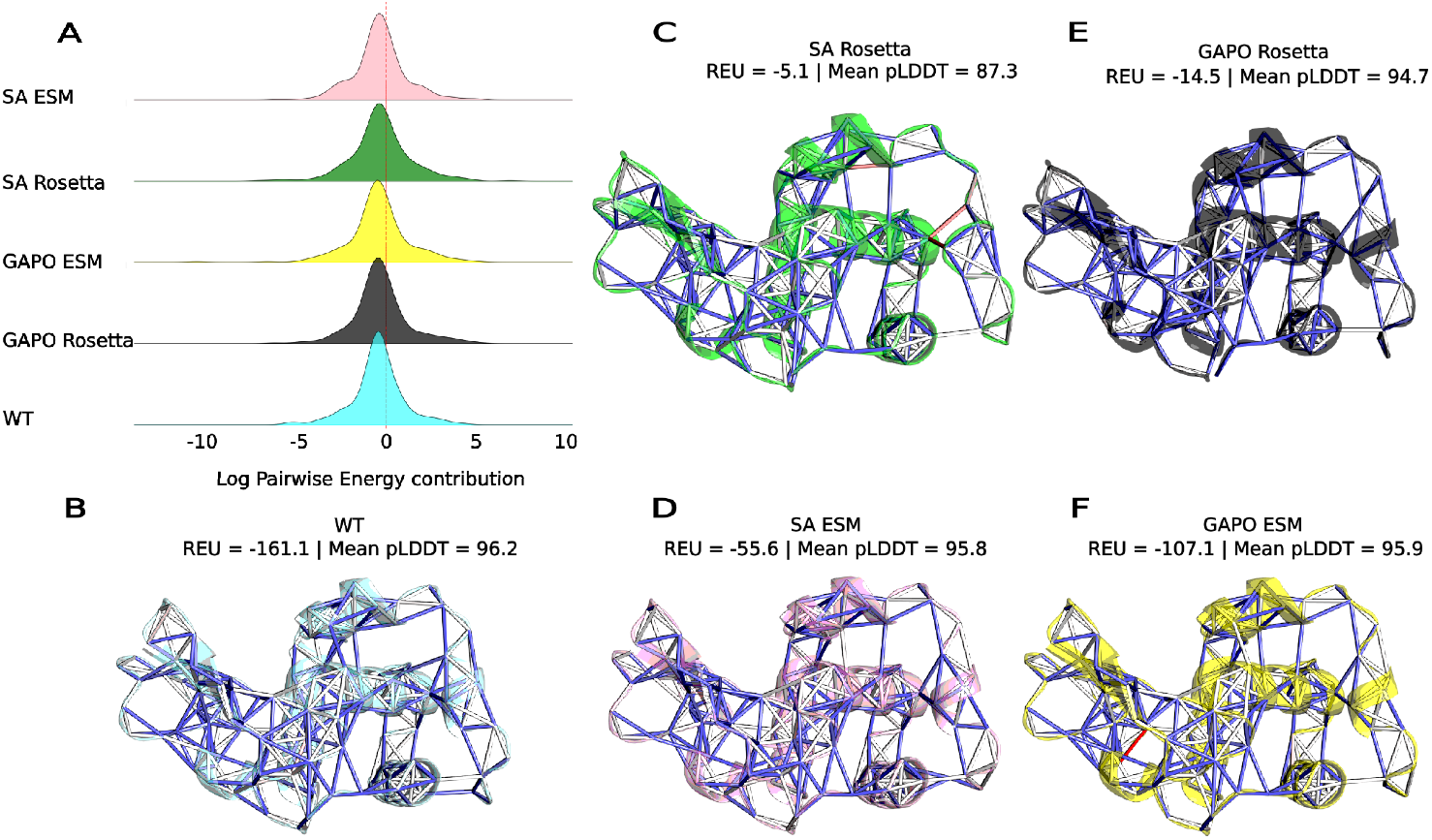
Comparative energetic and structural analysis of optimized lysozyme variants. (A) Distribution of pairwise non-bonded interaction energies for the wild-type (WT) and the top-performing variant from each condition, highlighting their overall energetic profiles. (B-F) Structural visualizations of the interaction energy networks. Cylinders connect the Cα atoms of interacting residues and are colored by interaction type: blue for favorable and red for unfavorable. To highlight the most critical energetic hotspots, only the top 5% of interactions by magnitude between residues within a 6 Å distance are shown. The structures displayed are: (B) WT, (C) Top SA/Rosetta, (D) Top SA/ESM, (E) GAPO/Rosetta, and (F) Top GAPO/ESM.

## Conclusions

Here, we introduced GAPO, a flexible genetic algorithm framework that provides a powerful alternative for protein sequence optimization and evaluated its utility through a proof-of-concept study. By optimizing lysozyme, GAPO significantly outperformed SA, a common optimization baseline, across all evaluated fitness functions tested. The framework’s primary strength lies in its flexibility, which affords broad applicability by allowing users to customize key components: the initial population strategy, the fitness function for optimization, and the choice of state-of-the-art PLMs to generate mutations.

Regarding initialization, both natural-sequence and ESM2-based seeding proved effective. However, the ESM2-seeded population explored more diverse regions of the sequence space, highlighting its utility for discovering novel variants. Conversely, the natural-sequence approach offered a more conservative trajectory, better preserving the original sequence identity and structural stability.

A notable finding from these initial tests is the effectiveness of mutations guided by the ESM2 in optimizing a force field-based metric, Rosetta’s REF15 energy function. This synergy suggests that PLMs capture abstract features of protein fitness that are complementary to traditional, physics-based energy functions, highlighting a promising avenue for the development of powerful hybrid methodologies.

The flexible architecture of GAPO is specifically designed to explore this frontier. While the results presented here serve as an initial validation using a simple, well-established test case, they underscore the framework’s potential. GAPO’s ability to combine distinct fitness functions directly enables the next step in this research: the concurrent optimization of a protein using both ESM2 and REF15. Such an approach would create a novel, composite fitness landscape that capitalizes on the complementary strengths of both data-driven and physics-based models.

Looking forward, the extensibility of GAPO will be crucial. As the field of protein engineering rapidly advances, new models and scoring functions can be readily integrated into the framework. This will facilitate not only more sophisticated sequence- and structure-driven design campaigns but also complex, multi-parameter optimization routines. The continued development of GAPO will therefore provide a valuable tool for pioneering and testing the next generation of hybrid protein design strategies.

## Acknowledgements

We gratefully acknowledge the use of computational resources provided by the RPT04A Bioinformatics Core Facility at Fiocruz, Rio de Janeiro. We gratefully acknowledge the financial support from Fundação Carlos Chagas Filho de Amparo à Pesquisa do Estado do Rio de Janeiro (Faperj) and Conselho Nacional de Desenvolvimento Científico e Tecnológico (CNPq). This study was financed in part by the Coordenação de Aperfeiçoamento de Pessoal de Nível Superior – Brasil (CAPES) – Finance Code 001.

## Code Availability

GAPO is available at https://github.com/izzetbiophysicist/GAPO.

## Author contributions

Sartori, J. implemented fitness functions, crossover modules, and interfaces with ESM2; managed all updates; ran and designed experiments; and wrote and reviewed the manuscript. Machado, L. A. conceived the idea; implemented GAPO’s core functionalities; contributed to the experimental design; and wrote and reviewed the manuscript. Krempser, E. advised on the implementation and theoretical background, and wrote and reviewed the manuscript. Guimarães, A. C. R. provided biological background for the experiments, supervised the tests, and wrote and reviewed the manuscript.

